# Cyclic Peptides Target CAPON and Modulate Cellular Responses under Alzheimer’s Disease-Relevant Stress

**DOI:** 10.64898/2026.05.10.724063

**Authors:** Ashraf Abdo, Shaoren Yuan, Katarzyna Kuncewicz, Jisong Mo, Hongliang Duan, Moustafa T. Gabr

## Abstract

CAPON (NOS1AP) is an adaptor protein involved in neuronal nitric oxide synthase (nNOS) signaling and has been implicated in Alzheimer’s disease (AD), excitotoxicity, and tau-associated neurodegeneration. Here, we report the identification of cyclic peptide ligands targeting CAPON using phage display screening of a disulfide-constrained peptide library. Phage enrichment, ELISA validation, microscale thermophoresis (MST), and biolayer interferometry (BLI) identified CAP1 as the lead peptide, exhibiting low micromolar binding affinity toward CAPON. Computational studies further supported stable CAPON–CAP1 interactions through complementary hydrophobic and electrostatic contacts. Functionally, CAP1 attenuated Aβ42-induced neuronal toxicity, suppressed NMDA-driven nitric oxide production, and reduced pathological tau phosphorylation in neuronal models under AD-relevant stress conditions. In addition, CAP1 demonstrated favorable preliminary pharmacokinetic properties, including good aqueous solubility, plasma stability, and measurable membrane permeability. Collectively, these findings establish the first cyclic peptide ligands targeting CAPON and identify CAP1 as a promising scaffold for modulation of CAPON-dependent neurodegenerative signaling.

## 1. Introduction

CAPON (NOS1AP) has emerged as an important regulator of multiple pathological processes, including neurodegenerative disorders, neurotoxicity, and cardiovascular disease. CAPON functions primarily as an adaptor protein for neuronal nitric oxide synthase (nNOS), where it regulates nitric oxide (NO) signaling within neurons.^1-3^ Beyond its role in NO signaling, CAPON has also been implicated in NMDA receptor-dependent pathways,^4^ contributing to the regulation of synaptic plasticity, neuronal development, and apoptosis.^5^ CAPON was initially identified as a carboxy-terminal PDZ ligand for nNOS through interactions mediated by its C-terminal PDZ-binding motif and the PDZ domain of nNOS.^6^

The PDZ domain of nNOS serves as a central hub for protein–protein interactions involving multiple neuronal signaling partners, including PSD-95 and CAPON, both of which compete for overlapping binding regions on nNOS.^7,8^ Under physiological conditions, PSD-95 recruits nNOS to NMDA receptor complexes, where calcium influx following glutamate stimulation activates nNOS and promotes tightly regulated NO production involved in learning, synaptic signaling, and neuronal plasticity.^2^ However, excessive glutamatergic stimulation results in pathological overactivation of nNOS, leading to elevated production of NO and peroxynitrite and ultimately contributing to neuronal injury and excitotoxicity.^9,10^

Within this signaling network, CAPON modulates nNOS function by competing with PSD-95 for association with the nNOS PDZ domain, thereby shifting nNOS toward alternative signaling complexes associated with reduced NO production and protection against excitotoxic stress.^11^ Importantly, this regulatory balance appears highly context dependent, as dysregulated or elevated CAPON expression may disrupt normal PSD-95-nNOS signaling and contribute to neuronal dysfunction.^9^

Disruption of the CAPON-nNOS signaling axis has been implicated in a broad range of neurological and psychiatric disorders, including Alzheimer’s disease (AD),^12^ Parkinson’s disease,^13^ Huntington’s disease,^14^ and Amyotrophic lateral sclerosis,^14^ as well as major depressive disorder,^15^ bipolar disorder,^5^ anxiety, and post-traumatic stress disorder (PTSD).^5^ Mechanistically, CAPON overexpression has been associated with abnormal Tau phosphorylation and formation of the Dexras1-nNOS-CAPON signaling complex, which promotes β-amyloid accumulation and accelerates AD-related pathology.^16,17^ These molecular alterations are subsequently linked to synaptic dysfunction and progressive neurodegeneration.^12^ In parallel, CAPON-dependent signaling has been shown to influence downstream MAPK pathways, including reduced ERK signaling and increased p38 activation, both of which are associated with neurotoxic responses.^11,18^

Additional studies in the hippocampal dentate gyrus further demonstrated that disruption of the nNOS-CAPON interaction alters responses to antidepressants such as fluoxetine. CAPON-mediated modulation of nNOS signaling affects phosphorylation levels of ERK, CREB, and BDNF, correlating with behavioral phenotypes associated with anxiety and depression.^11,18^ Collectively, these findings support the CAPON-nNOS signaling axis as an attractive yet still underexplored therapeutic target for neurological and neuropsychiatric disorders.

Peptide therapeutics have increasingly emerged as an intermediate modality bridging traditional small molecules and biologics. In particular, cyclic peptides offer several advantages over their linear counterparts, including enhanced conformational stability, improved resistance to proteolytic degradation, and increased affinity toward protein surfaces.^19^ Furthermore, the constrained cyclic scaffold enables efficient engagement of shallow or extended binding interfaces commonly associated with protein–protein interactions (PPIs).^20^ As a result, cyclic peptides represent a promising strategy for targeting proteins and interaction surfaces that have traditionally been considered difficult to drug using conventional small molecules.

In this study, we employed phage display screening of a disulfide-constrained cyclic peptide library to identify ligands capable of directly targeting CAPON. Selected peptides were subsequently validated using orthogonal biophysical approaches, including microscale thermophoresis (MST) and biolayer interferometry (BLI), to confirm binding and characterize binding affinity. Functional evaluation in AD-relevant neuronal models demonstrated that the lead peptide, CAP1, attenuates Aβ42-induced neurotoxicity, suppresses NMDA-driven nitric oxide signaling, and reduces pathological tau phosphorylation. Collectively, these findings establish the first cyclic peptide ligands targeting CAPON and identify CAP1 as a promising scaffold for modulation of CAPON-dependent neurodegenerative signaling.

## 2. Results and discussion

### Phage display reveals cyclic peptide binders targeting CAPON

To identify cyclic peptide ligands capable of engaging CAPON, we performed phage display selection using a disulfide-constrained MC9 cyclic peptide library based on a CX_9_C scaffold (Figure 1A). Recombinant human CAPON (NOS1AP) protein was used as the target for iterative rounds of biopanning aimed at enriching peptide sequences capable of recognizing the CAPON surface. Four rounds of selection, including binding, washing, elution, and amplification, were performed to progressively enrich CAPON-binding phage populations.

**Figure 1.**
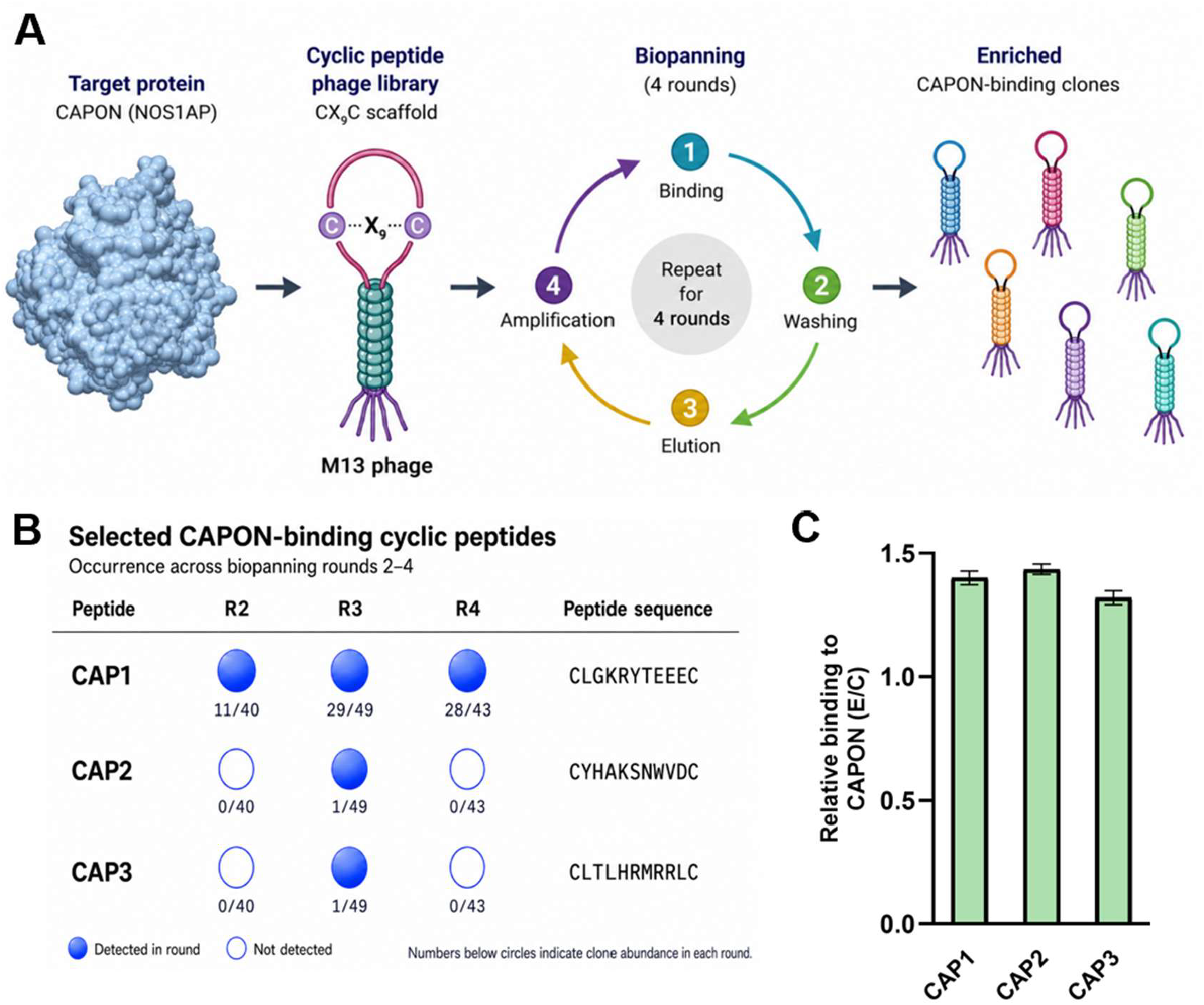
Identification and validation of CAPON-binding cyclic peptides by phage display. **(A)** Schematic illustration of the phage display workflow. Recombinant CAPON (NOS1AP) protein was used as the target for selection from a cysteine-constrained cyclic peptide M13 phage library. Four rounds of biopanning, including binding, washing, elution, and amplification, were performed to enrich CAPON-binding clones. **(B)** Top enriched cyclic peptide sequences identified across rounds 2-4 of biopanning. Filled circles indicate detection of a given sequence in the corresponding round, whereas open circles indicate absence. CAP1 (CLGKRYTEEEC) emerged as the dominant enriched sequence across successive rounds of selection. **(C)** Phage ELISA validation of selected CAPON-binding peptides. Binding signals are presented as the ratio of signal obtained in CAPON-coated wells relative to control wells lacking protein (E/C). Values above 1 indicate preferential binding to CAPON. Data are shown as mean ± SEM (n = 3).

Analysis of the output libraries revealed clear enrichment during the selection process, particularly during rounds 2 and 3, consistent with preferential amplification of CAPON-binding clones (Figure 1A). Beginning in round 2, individual phage clones were isolated and sequenced to identify enriched cyclic peptide motifs. Among the identified sequences, CLGKRYTEEEC emerged as the dominant enriched peptide across successive rounds of selection, accounting for 11/40 clones in round 2, 29/49 clones in round 3, and 28/43 clones in round 4 (Figure 1B). In contrast, CYHAKSNWVDC and CLTLHRMRRLC were detected at lower frequency, each appearing once during round 3 sequencing. Despite their lower enrichment frequency, these sequences were retained for further evaluation based on their sequence diversity and distinct physicochemical properties.

Based on enrichment behavior and sequence diversity, three peptides were selected for downstream characterization and designated CAP1 (CLGKRYTEEEC), CAP2 (CYHAKSNWVDC), and CAP3 (CLTLHRMRRLC). Notably, all three peptides preserved the cysteine-constrained cyclic scaffold while exhibiting substantial variation within the internal residues, suggesting that CAPON recognition may occur through multiple peptide interaction modes rather than convergence toward a single consensus binding motif. This observation is consistent with the adaptor-like and protein-protein interaction-driven nature of CAPON.

To further evaluate peptide binding, representative phage clones were analyzed by phage ELISA against immobilized CAPON protein (Figure 1C). All three selected peptides exhibited preferential binding to CAPON relative to control wells lacking protein, supporting selective target recognition. Among the evaluated sequences, CAP1 demonstrated the strongest enrichment profile during biopanning together with robust ELISA binding signals, supporting its prioritization for subsequent orthogonal biophysical characterization. Collectively, these findings demonstrate that cyclic peptide phage display can successfully identify ligands targeting CAPON and establish a focused set of candidates for downstream functional studies.

### Biophysical validation for CAPON binding

Prior to biophysical characterization, all cyclic peptides were evaluated for aqueous solubility. No visible precipitation or aggregation was observed for any of the tested peptides under the assay conditions, supporting that the measured binding responses were not driven by solubility-related artifacts.

To evaluate direct binding to CAPON, the selected cyclic peptides were analyzed using MST. As shown in Figure 2, CAP1, CAP2, and CAP3 produced distinct concentration-dependent binding curves, consistent with direct interaction with CAPON. Dose-dependent MST experiments were subsequently performed to determine the apparent dissociation constants (Kd) for the peptides exhibiting measurable binding. Nonlinear regression analysis of the binding curves revealed apparent Kd values of 2 ± 3 µM for CAP1 (Figure 2A), 19 ± 15 µM for CAP2 (Figure 2B), and 194 ± 50 µM for CAP3 (Figure 2C). Among the evaluated peptides, CAP1 displayed the strongest binding affinity toward CAPON, consistent with its dominant enrichment profile during phage display selection. CAP2 also demonstrated low micromolar binding, whereas CAP3 exhibited substantially weaker affinity.

**Figure 2.**
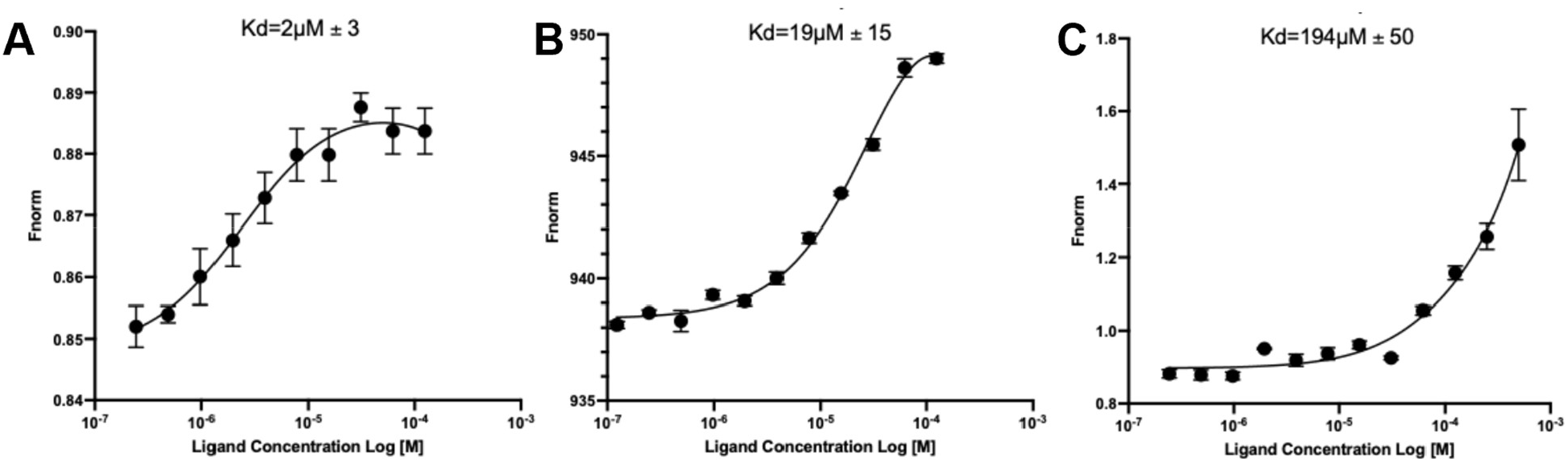
Results for Monolith binding studies. MST-based characterization of CAPON-binding cyclic peptides. Dose-dependent MST binding curves for CAP1 **(A)**, CAP2 **(B)**, and CAP3 **(C)** against recombinant CAPON protein. Normalized fluorescence (Fnorm) is plotted as a function of peptide concentration on a logarithmic scale. Binding curves were fitted using nonlinear regression analysis to determine apparent dissociation constants (Kd). CAP1 exhibited the strongest binding affinity (Kd = 2 ± 3 µM), followed by CAP2 (Kd = 19 ± 15 µM) and CAP3 (Kd = 194 ± 50 µM). Data points represent mean ± SEM from independent measurements.

Importantly, the peptides validated by MST directly corresponded to sequences identified during phage display enrichment, providing orthogonal confirmation of the selection outcome. Collectively, these findings establish cyclic peptide binding to CAPON and identify CAP1 as a promising lead scaffold for further computational, pharmacokinetic (PK), and functional characterization.

To further validate peptide binding using an orthogonal biophysical approach, the most potent peptides identified from MST analysis, CAP1 and CAP2, were subsequently evaluated by BLI. In these experiments, recombinant CAPON protein was immobilized on the biosensor surface, and peptide binding was assessed across increasing peptide concentrations to characterize steady-state binding behavior.

As shown in Figure 3, both CAP1 and CAP2 produced concentration-dependent BLI responses consistent with direct interaction with CAPON. CAP1 displayed rapid association kinetics together with strong saturation behavior at low micromolar concentrations (Figure 3A). Steady-state fitting using a one-site specific binding model yielded a Kd value of 1.3 ± 0.2 µM, supporting high-affinity interaction between CAP1 and CAPON. In contrast, CAP2 required substantially higher concentrations to achieve saturation and exhibited weaker overall binding affinity, with an apparent KD of 15.3 ± 2.8 µM (Figure 3B). Importantly, the relative binding hierarchy observed by BLI was fully consistent with the MST results, where CAP1 similarly emerged as the strongest binder among the evaluated peptides.

**Figure 3.**
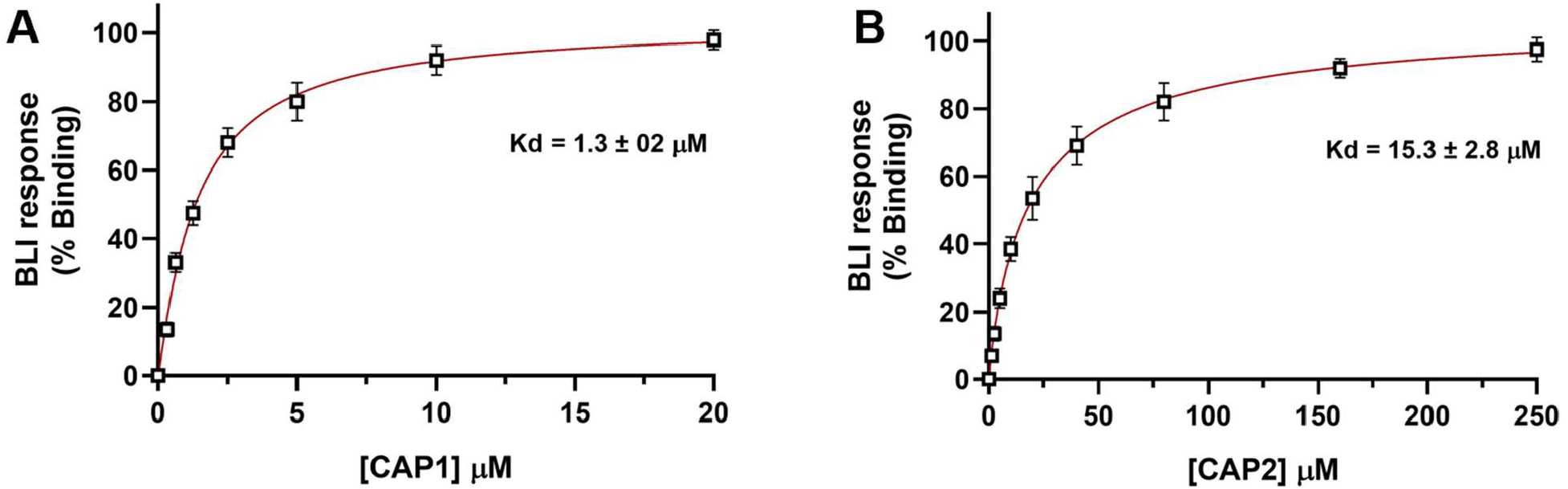
Steady-state BLI analysis of CAPON-binding cyclic peptides. Steady-state biolayer interferometry (BLI) binding analysis of CAP1 **(A)** and CAP2 **(B)** interacting with immobilized CAPON protein. BLI responses were measured across increasing peptide concentrations and independently normalized to the maximal binding response for each peptide. Binding curves were fitted using a one-site specific binding model to estimate apparent dissociation constants (Kd). CAP1 exhibited stronger binding affinity toward CAPON (Kd = 1.3 ± 0.2 µM) relative to CAP2 (Kd = 15.3 ± 2.8 µM). Data points represent mean ± SEM from independent measurements.

Notably, both peptides demonstrated saturable binding profiles characteristic of specific target engagement rather than nonspecific surface interactions. The stronger binding affinity of CAP1 is also consistent with its dominant enrichment behavior during phage display selection, suggesting that iterative biopanning successfully enriched for higher-affinity CAPON-binding sequences. Collectively, these findings further establish CAP1 as the lead cyclic peptide scaffold identified in this study and support its prioritization for characterization in AD-relevant cellular models.

### CAP1 modulates CAPON-dependent neurodegenerative signaling pathways

To determine whether CAPON modulation by CAP1 confers broader neuroprotective activity, we evaluated neuronal viability following exposure to Aβ42 oligomers. Pretreatment with CAP1 produced a concentration-dependent reduction in LDH release relative to Aβ42-treated controls, consistent with attenuation of amyloid-associated neurotoxicity (Figure 4A). CAP1 alone did not significantly affect basal cytotoxicity, further supporting the specificity of the observed protective effects.

**Figure 4.**
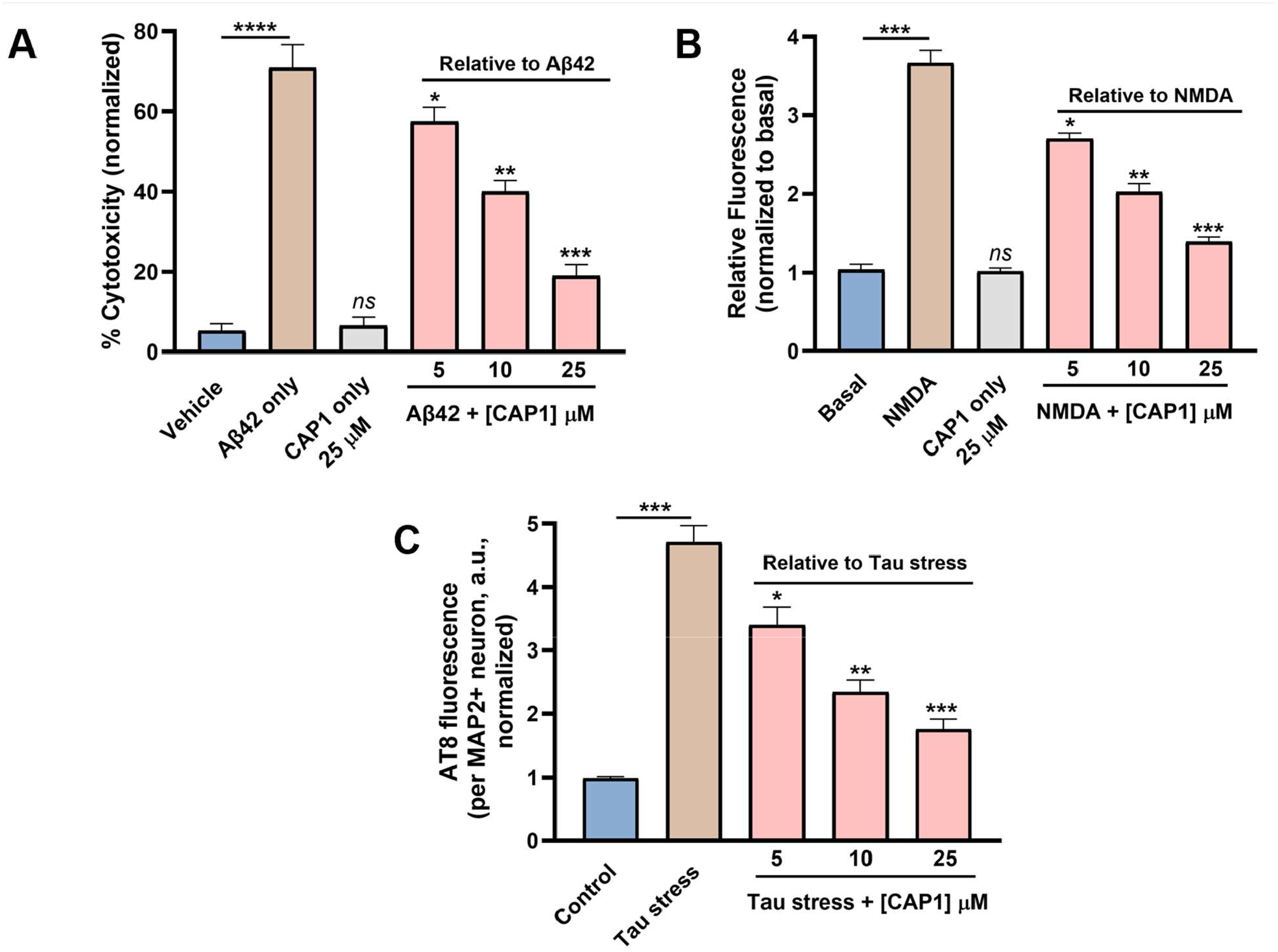
CAP1 modulates CAPON-dependent neurodegenerative signaling pathways in neuronal models. **(A)** CAP1 protects primary cortical neurons from Aβ42-induced cytotoxicity. Neurons were pretreated with CAP1 (5, 10, or 25 µM) prior to exposure to Aβ42 oligomers, and cytotoxicity was quantified by LDH release assay. CAP1 produced a concentration-dependent reduction in Aβ42-induced neuronal toxicity relative to Aβ42-treated controls. **(B)** CAP1 suppresses pathological NMDA-driven nitric oxide signaling. Intracellular nitric oxide production was measured using DAF-FM DA fluorescence following NMDA stimulation in the presence or absence of CAP1. CAP1 attenuated NMDA-induced NO production in a concentration-dependent manner while showing no significant effect alone. **(C)** CAP1 reduces pathological tau phosphorylation in neuronal cells. Neurons subjected to tau-stress conditions were treated with CAP1, and tau pathology was assessed by AT8 immunofluorescence. Quantitative analysis demonstrated a concentration-dependent reduction in AT8 fluorescence intensity per MAP2^+^ neuron relative to tau-stressed controls. Data represent mean ± SEM from independent measurements. *p < 0.05, **p < 0.01, ***p < 0.001, ****p < 0.0001, and ns denotes nonsignificant relative to the indicated control group.

To further determine whether the lead cyclic peptide CAP1 modulates CAPON-dependent signaling downstream of NMDA receptor activation, we first evaluated intracellular nitric oxide (NO) production using the fluorescent probe DAF-FM DA (4-amino-5-methylamino-2′,7′-difluorofluorescein diacetate). Acute NMDA stimulation produced a pronounced increase in intracellular NO levels, consistent with activation of nNOS-dependent nitrosative signaling pathways. Pretreatment with CAP1 significantly attenuated NMDA-induced NO production in a concentration-dependent manner, with partial suppression observed at 5 µM and progressively stronger inhibition observed at 10 and 25 µM (Figure 4B). Importantly, CAP1 alone did not significantly alter basal NO production, suggesting that modulation of the CAPON–nNOS axis preferentially suppresses pathological NMDA-driven signaling without globally impairing physiological neuronal activity. Because nitric oxide production represents a proximal downstream readout of nNOS activation, these findings support the ability of CAP1 to functionally interfere with CAPON-dependent signaling associated with excitotoxic stress.

Given the established role of CAPON signaling in regulating tau-associated neurodegenerative pathways, we next examined whether CAP1 modulates pathological tau phosphorylation in neuronal cells. Under tau-stress conditions, neuronal cultures exhibited a marked increase in AT8 immunoreactivity, consistent with enhanced phosphorylation of tau at disease-relevant epitopes. Treatment with CAP1 resulted in a concentration-dependent reduction in AT8 fluorescence intensity per MAP2^+^ neuron (Figure 4C). Notably, CAP1 reduced pathological tau phosphorylation without significantly affecting neuronal morphology or MAP2 labeling, indicating that the observed effects were not attributable to nonspecific neuronal toxicity. These findings demonstrate that CAP1 suppresses pathological tau-associated signaling under disease-relevant stress conditions. Collectively, these findings demonstrate that CAP1 exerts convergent protective effects across multiple pathological pathways associated with AD, including amyloid toxicity, nitrosative stress, and tau dysregulation. By suppressing NMDA-dependent nitric oxide signaling and attenuating pathological tau phosphorylation, CAP1 modulates downstream processes closely linked to excitotoxicity and neurodegeneration.

### Computational analysis of CAPON-peptide interactions

Molecular dynamics (MD) simulations were performed to evaluate the binding stability of CAPON with CAP1 and CAP2. As shown by the RMSD trajectories (Figure 5), the two peptides displayed distinct dynamic behaviors during the simulation. The CAPON–CAP1 complex exhibited RMSD fluctuations throughout the simulation, suggesting that this system underwent pronounced conformational rearrangement before reaching a stable bound conformation. In contrast, the CAPON–CAP2 complex, the RMSD increased during the early stage and then remained relatively stable with moderate fluctuations during the later stage, indicating that the complex progressively converged to a stable bound conformation.

**Figure 5.**
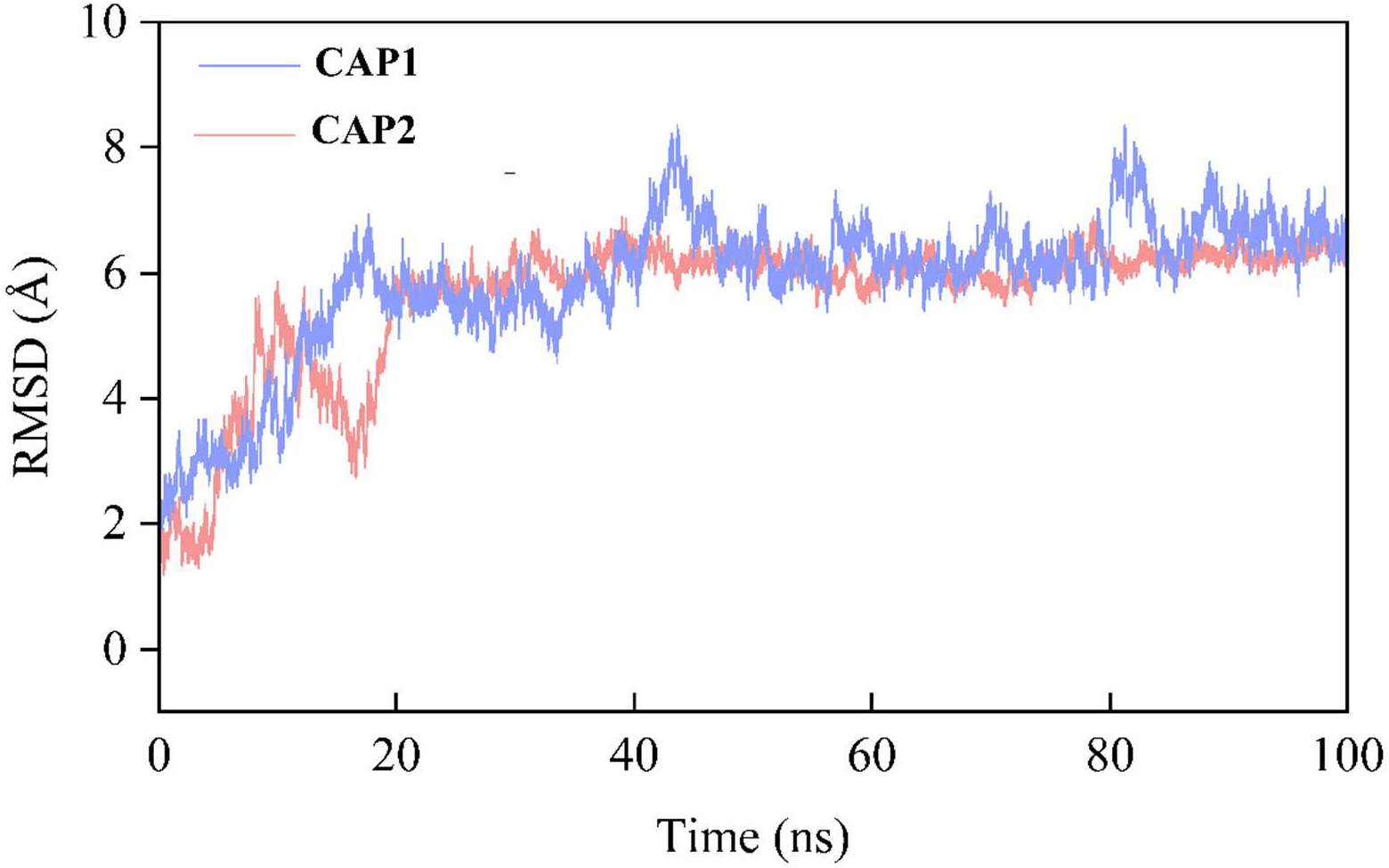
Root mean square deviation (RMSD) profiles of the CAPON–CAP1 and CAPON– CAP2 complexes during 100 ns molecular dynamics simulations.

Representative structures extracted from the last 50 ns of each trajectory showed that the two peptides stabilized distinct binding interfaces on the CAPON surface (Figure 6). In the CAPON– CAP1 complex (Figure 6A), the interface is substantially broader, engaging Tyr114, Phe117, Tyr118, Ser120, Phe165, Glu166, Leu173, Gln177, Lys195, Asn197, Trp198, and Asp200. Beyond hydrophobic contacts from Phe117, Leu173, and Trp198, and polar interactions from Tyr114, Ser120, Gln177, and Asn197, CAP1 uniquely engages three charged CAPON residues (Glu166, Lys195, and Asp200) introducing a denser network of electrostatic contacts absent from the CAP2 interface. In the CAPON–CAP2 complex (Figure 6B), the interface is anchored by hydrophobic contacts with Ile116, Phe165, and Leu192, supplemented by hydrogen bonds through Tyr118, Ser172, Thr176, and Tyr196, and an electrostatic interaction with Glu198, forming a compact, hydrophobically dominated contact region. Tyr118 and Phe165 are shared by both complexes, constituting a conserved aromatic anchor, while the divergent peripheral networks reflect the distinct amino acid compositions of the two peptides and their differential complementarity to the CAPON surface. Notably, the conformational differences observed in the CAPON binding region between the two complexes may be attributed to the intrinsic flexibility of local loop segments, which were selectively remodeled by the distinct physicochemical properties of each peptide during MD relaxation.

**Figure 6.**
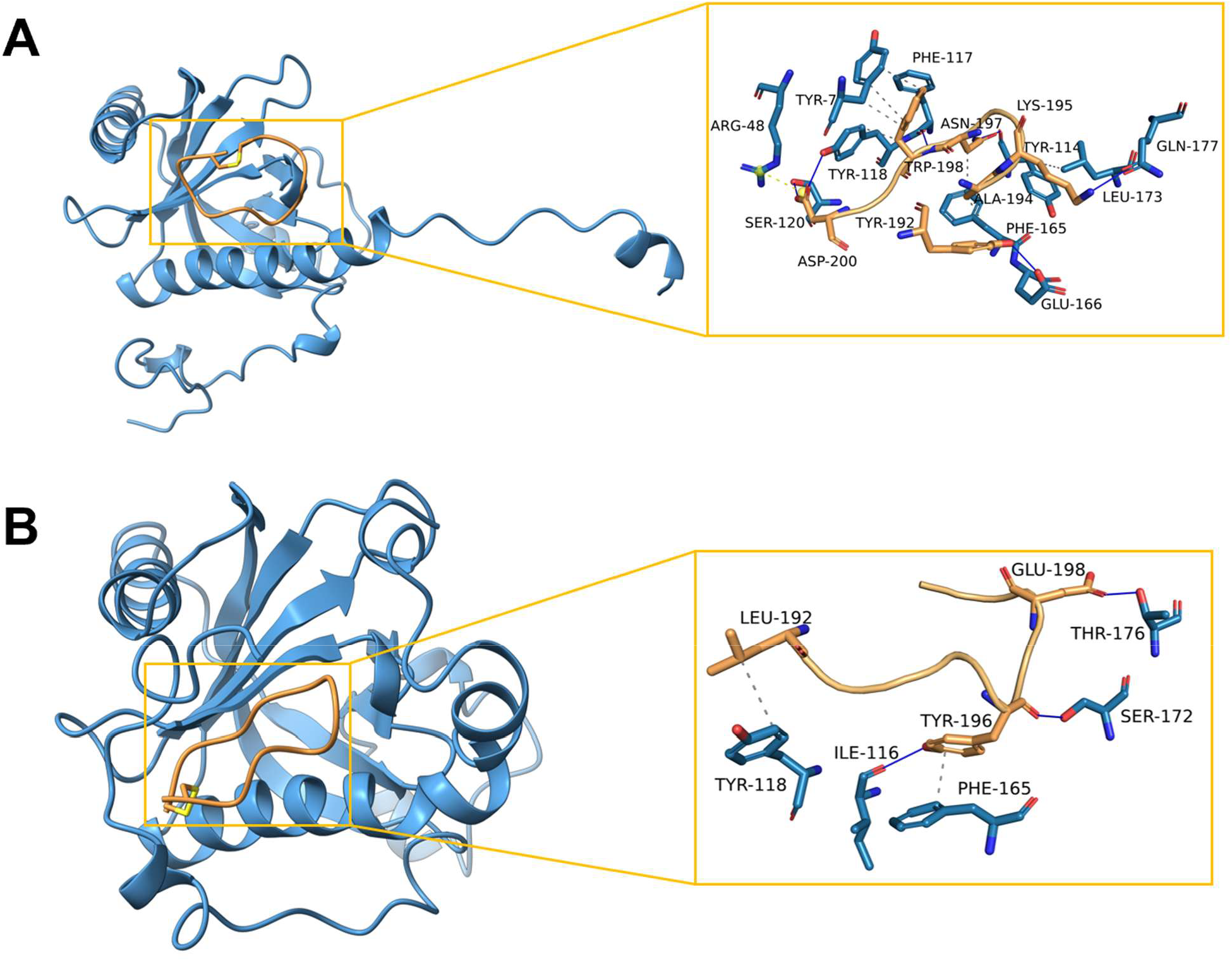
Structural comparison of the binding modes of CAP1 **(A)**, and CAP2 **(B)** on the CAPON surface. In the binding model, hydrophobic interactions are indicated by grey dashed lines, hydrogens bonds by navy blue solid lines, salt bridges by yellow dashed lines.

### Preliminary PK characterization of CAP1

Following identification of CAP1 as the lead CAPON-binding cyclic peptide based on phage display enrichment, MST, BLI, and functional cellular assays, we next evaluated its preliminary in vitro PK properties to assess developability characteristics relevant to peptide-based therapeutics. Because cyclic peptide scaffolds can exhibit improved metabolic stability and proteolytic resistance relative to linear peptides, we sought to determine whether CAP1 retained favorable stability and permeability properties under physiologically relevant conditions.

As summarized in Table 1, CAP1 demonstrated measurable stability in both simulated gastric and intestinal fluids, exhibiting half-lives of 1.12 h and 6.84 h, respectively. The substantially improved stability observed under intestinal conditions is consistent with the conformationally constrained cyclic scaffold and suggests partial resistance to proteolytic degradation. In human plasma, CAP1 retained 81.3% integrity after 1 h incubation, further supporting favorable plasma stability for an early-stage cyclic peptide lead.

**Table 1.** *In vitro* PK profile of **CAP1**.

| Test | CAP1 |
| --- | --- |
| Stability in simulated fluids ($t_{1/2}$, h) | |
| Gastric (pH 1.6) | 1.12 |
| Intestinal (pH 6.5) | 6.84 |
| Stability in human plasma (% remaining at 1 h) | 81.3 |
| Aqueous solubility at pH 7.4 (μM) | 148.7 |
| PPB (% bound, fu) | 22.4 |
| $\text{Log}D_{7.4}^a$ | 0.67 |
| CACO cell permeability<sup>b</sup> |  |
| $P_{\text{app}, A \rightarrow B}$ ($10^{-6}$ cm/s) | 1.46 |
| $P_{\text{app}, B \rightarrow A}$ ($10^{-6}$ cm/s) | 2.11 |
| Stability in rat liver microsomes ($t_{1/2}$, h) | 3.42 |

<sup>a</sup>Determined by miniaturized shake-flask procedure at pH7.4.
<sup>b</sup>Permeation through a Caco-2 cell monolayer was assessed in the absorptive apical to basolateral (a→b) and secretory basolateral to apical (b→a).

CAP1 also displayed good aqueous solubility at physiological pH (148.7 µM at pH 7.4), supporting its suitability for downstream biochemical and cellular studies. Measurement of plasma protein binding indicated moderate protein association, with a free fraction consistent with appreciable levels of unbound peptide remaining available for target engagement. In parallel, CAP1 exhibited a LogD7.4 value of 0.67, suggesting a balanced physicochemical profile compatible with aqueous solubility while still maintaining moderate hydrophobic character.

To further evaluate membrane permeability, CAP1 was analyzed using the CACO-2 permeability assay. CAP1 demonstrated measurable bidirectional permeability with an apparent permeability coefficient (Papp) of 1.46 × 10^−6^ cm/s in the apical-to-basolateral direction and 2.11 × 10^−6^ cm/s in the reverse direction. Although permeability remained within the expected range for cyclic peptides, these findings nevertheless suggest that CAP1 retains partial membrane transport capability despite its peptide-based structure.

Finally, metabolic stability studies using rat liver microsomes demonstrated a half-life of 3.42 h, indicating moderate resistance to microsomal degradation. Collectively, these findings demonstrate that CAP1 possesses a favorable early-stage PK profile characterized by good aqueous solubility, measurable plasma and microsomal stability, and detectable membrane permeability. Together with its orthogonal biophysical validation and functional activity in AD-relevant cellular models, these data further support CAP1 as a promising cyclic peptide scaffold for continued optimization targeting the CAPON–nNOS signaling axis.

In summary, this study reports the identification and characterization of the first cyclic peptide ligands targeting CAPON, a therapeutically relevant adaptor protein implicated in excitotoxicity and AD-related neurodegeneration. Using phage display screening of a disulfide-constrained cyclic peptide library, followed by orthogonal biophysical validation by MST and BLI, we identified CAP1 as a lead CAPON-binding peptide with low micromolar affinity. Computational studies further supported stable CAPON–CAP1 interactions and revealed a binding interface involving complementary hydrophobic, polar, and electrostatic contacts.

Importantly, CAP1 demonstrated functional activity across multiple AD-relevant cellular pathways, including suppression of NMDA-driven nitric oxide signaling, attenuation of pathological tau phosphorylation, and protection against Aβ42-induced neuronal toxicity. In parallel, CAP1 exhibited favorable preliminary PK and physicochemical properties, including good aqueous solubility, measurable plasma and microsomal stability, and detectable membrane permeability.

Collectively, these findings establish cyclic peptides as a viable modality for targeting the CAPON–nNOS signaling axis and provide a foundation for future optimization of CAPON-directed ligands for neurodegenerative disease applications. Beyond AD, the broader involvement of CAPON in neurological and psychiatric disorders further highlights the therapeutic potential of targeting this adaptor protein using peptide-based approaches.

## 3. Materials and methods

### Phage display selection of CAPON-binding cyclic peptides

A disulfide-constrained phage-displayed cyclic peptide library based on a CX_9_C scaffold was used to identify ligands capable of engaging human CAPON. Recombinant human CAPON (Origene, Catalog# TP760978) was immobilized on a solid support, and the phage library was subjected to iterative rounds of biopanning against the target protein. After each round, nonbinding phages were removed by washing, whereas bound phages were recovered by elution and amplified in *E. coli* for use in the subsequent round of selection. A total of four rounds of biopanning were performed, with enrichment of CAPON-binding phage monitored across rounds.

Following the final selection round, phage pools were analyzed by next-generation sequencing to identify enriched peptide sequences. Sequence analysis revealed recurrent CX_9_C-containing clones that increased in frequency over the course of selection, consistent with preferential enrichment of CAPON-binding peptides. Representative enriched clones were designated and prioritized for downstream synthesis and functional characterization.

To assess clone-level binding, selected phages were evaluated by phage ELISA against immobilized recombinant CAPON. Briefly, target-coated wells were incubated with individual phage clones, washed to remove unbound material, and bound phage were detected using an anti-M13 horseradish peroxidase-conjugated antibody and colorimetric substrate. Signal intensities were compared with background controls to identify clones exhibiting specific binding to CAPON. Clones showing reproducible enrichment and positive phage ELISA signals were advanced for chemical synthesis and orthogonal biophysical validation.

### Peptide synthesis and characterization

Peptide synthesis was carried out by solid-phase peptide synthesis (SPPS) via the Fmoc/tBu strategy using an automated microwave peptide synthesizer, Liberty BLUE (CEM, Matthews, NC, USA). 424 mg Rink Amide ProTide resin (0.59 mmol/g; CEM, Matthews, NC, USA) was used to synthesize each peptide. During the synthesis, each amino acid was coupled twice using a 5-fold excess relative to the resin loading. The following solutions were used during the synthesis: OxymaPure/DIC as coupling reagents, 20% piperidine in DMF for Fmoc deprotection, and DMF for washing between the deprotection and coupling steps.

Synthesized peptides were cleaved from the resin using a mixture containing 88% TFA, 5% H_2_O, 5% phenol, and 2% TIPS (Reagent B; 10 mL of solution was used for 424 mg of resin). Crude peptides were precipitated with cold diethyl ether, decanted, and lyophilized. Initial purification was performed if low purity of the crude uncyclized peptide was observed by LC–MS. For purification, the peptides were dissolved in water with the addition of a 10-fold molar excess of dithiothreitol (DTT) relative to free sulfhydryl groups and incubated for 30 minutes at 60 °C. Linear peptides were purified by reversed-phase HPLC using an XBridge Prep C18 column (19 × 150 mm, 5 μm, Waters, MA, USA) at a flow rate of 20 mL/min with a 10 min run using 5–95% acetonitrile in water containing 0.1% formic acid. The purity and mass spectra of the final products were analyzed using an ACQUITY HPLC system equipped with an SQ Detector 2 and an XBridge C18 analytical column (4.6 × 150 mm, 5 μm, Waters, MA, USA), employing a linear gradient from 5% to 95% acetonitrile in water containing 0.1% formic acid over 10 minutes.

Oxidation of the peptides was performed using compressed air. The peptide was dissolved in H_2_O and methanol (1:9, v/v) at a concentration of approximately 60 mg/L, and the pH was adjusted and maintained between 8 and 9 using ammonia. The solution was stirred at room temperature for 3–5 days while compressed air was bubbled through the solution. After this time, the solvents were evaporated, and the peptides were lyophilized. Reaction progress was monitored using an analytical ACQUITY HPLC system equipped with an SQ Detector 2.

After this process, the peptides were purified again using the same ACQUITY HPLC system on an XBridge Prep C18 column (19 × 150 mm, 5 μm, Waters, MA, USA). A linear gradient of 5– 60% acetonitrile in water containing 0.1% formic acid over 10 min was used.

### Monolith binding assay

Peptides were evaluated in a concentration-dependent MST assay using Monolith X (from NanoTemper Technologies). The assay was performed as we previously reported.^21^ Briefly, twelve-point serial dilutions were prepared, spanning a starting concentration of 250 µM down to the low-nanomolar range, and mixed with labeled CAPON-His protein (Origene, Catalog# TP760978) while maintaining a final DMSO concentration of 2%. After a 30-minute equilibration step at room temperature, samples were loaded into standard MST capillaries and measured on a Monolith NT.115 instrument, using medium-to-high infrared laser power and 60–80% LED excitation in the red detection channel. Dissociation constants (Kd) were calculated using the MO.Affinity Analysis package (NanoTemper Technologies) as previously reported.^21^

### Biolayer interferometry (BLI) binding assay

BLI experiments were performed using an Octet RED96 system (Sartorius/ForteBio) to orthogonally validate binding of cyclic peptides to CAPON. All measurements were conducted at 25 °C with orbital shaking at 1000 rpm in black 96-well microplates containing 200 µL per well.

Recombinant human CAPON (Origene, Catalog# TP760978) protein was diluted in PBS and immobilized onto Ni-NTA biosensors pre-equilibrated in assay buffer. Prior to loading, biosensors were hydrated in assay buffer consisting of PBS supplemented with 0.01% Tween-20 and 0.1% bovine serum albumin (BSA) for at least 10 min. CAPON loading was performed for 300 s to achieve a stable immobilization signal, followed by a baseline equilibration step in assay buffer for 120 s.

For binding analysis, cyclic peptides CAP1 and CAP2 were prepared as serial dilutions in assay buffer. CAP1 was tested at concentrations ranging from 0.3125 to 20 µM, whereas CAP2 was tested from 0.625 to 250 µM. Association was monitored by transferring CAPON-loaded biosensors into peptide-containing wells for 300 s, followed by dissociation in peptide-free assay buffer for 600 s. Reference biosensors lacking immobilized protein together with buffer-only controls were included in each experiment for background subtraction and correction of nonspecific signal drift.

Raw sensorgrams were processed using ForteBio data analysis software. Signals from reference sensors and buffer controls were subtracted from experimental traces prior to analysis. Steady-state response values were extracted from the equilibrium region of the association phase and independently normalized to the maximal response obtained for each peptide. Binding curves were generated by plotting normalized BLI response as a function of peptide concentration and fitted using a one-site specific binding nonlinear regression model in GraphPad Prism 10. Apparent dissociation constants (Kd) are reported as mean ± SEM from independent measurements.

### Computational studies

In this study, molecular dynamics (MD) simulations were performed at 300 K for three systems: CAPON–CAP1 complexes, and CAPON–CAP2 complexes, using the AMBER20 software package^22^ combined with the ff14SB force field.^23^ The initial structures of the CAPON–CAP1 and CAPON–CAP2 complexes were generated by AlphaFold3 prediction.^24^ Before simulation, non-interfacial residues were removed so that the calculations were focused on the CAPON–peptide binding region. All solutes were placed in a truncated octahedral water box and solvated using the TIP3P water model,^25^ with the minimum distance from the solute surface to the box boundary set to 15.0 Å. Prior to the formal MD simulations, a two-stage energy minimization process was carried out on the systems to eliminate unfavorable steric clashes: (1) first, under positional constraints applied to the solutes, 5000 steps of steepest descent (SD) and 5000 steps of conjugate gradient (CG) energy optimization were performed; (2) subsequently, all constraints were removed, and an unconstrained energy minimization consisting of another 5000 steps of SD and 5000 steps of CG was executed.

After energy optimization, the systems underwent a heating process under positional constraints, during which the temperature was gradually increased from 0 K to 300 K. Subsequently, a 100 ns production simulation under the NPT (constant number of particles, pressure, and temperature) ensemble was conducted at 300 K and 1.0 bar. During the simulation, the temperature was controlled using the Nose-Hoover chain method,^26^ and the pressure was regulated by the Berendsen method.^27^ The SHAKE algorithm^28^ was employed to constrain all covalent bonds involving hydrogen atoms. The cutoff radius for non-bonded interactions was set to 10.0 Å, and long-range electrostatic interactions were calculated using the smooth Particle Mesh Ewald (PME) algorithm.^29^ The integration time step was set to 2 fs, and trajectory conformations were saved every 10 ps. Consequently, a total of 10,000 conformational frames were collected for each system for subsequent structural and interaction analyses.

### Evaluation of PK and physicochemical properties

These experiments were conducted following our previously reported methods.^30^

### Aβ_**42**_-Induced Neurotoxicity and Neuroprotection Assay

Primary cortical neurons (from E18 rat embryos) were cultured on poly-D-lysine–coated plates in Neurobasal medium supplemented with B27 and GlutaMAX. Neurons were used between DIV7-9. For Aβ_42_ oligomer preparation, Aβ_42_ peptide was dissolved in HFIP, dried to a film, resuspended in DMSO, diluted in phenol-red–free F12 medium, and incubated at 4 °C for 24 h. Neurons were pretreated with **CAP1** (5, 10, and 25 μM) or vehicle for 1 h prior to exposure to Aβ_42_ oligomers.

Following 24 h incubation, neuronal cytotoxicity was quantified using an LDH release assay. Maximum LDH release was determined using lysis buffer-treated wells, and background was subtracted using no-cell controls. In selected experiments, neuronal viability was confirmed using complementary live/dead staining or ATP-based viability assays.

### NMDA-Induced Nitrosative Stress Signaling

Primary cortical neurons (DIV8) were pretreated with **CAP1** (5, 10, and 25 μM) or vehicle for 1 h, followed by brief NMDA stimulation (50 µM, 10 min) to activate nNOS-dependent signaling. Cells were then washed and processed immediately. Nitric oxide (NO) production was quantified using the fluorescent probe DAF-FM DA, loaded into neurons prior to NMDA stimulation. Fluorescence intensity was measured by a Tecan Spark plate reader.

### Modulation of Tau Pathology in Neuronal Cells

Primary cortical neurons were cultured as described above and used between DIV7-10. To induce tau pathology (tau-stress), neurons were treated with okadaic acid (OA, 25 nM) for 24 h, to promote tau hyperphosphorylation through inhibition of protein phosphatases. Neurons were treated with **CAP1** (5, 10, and 25 μM) or vehicle control for 24 h.

Following treatment, cells were fixed with 4% paraformaldehyde, permeabilized, and blocked using standard conditions. Neurons were incubated with an antibody against phosphorylated tau (AT8; Ser202/Thr205) together with an antibody against MAP2 to label neuronal cell bodies and neurites. Nuclei were counterstained with DAPI. Fluorescent secondary antibodies were applied, and cells were imaged using a Tecan Spark. MAP2-positive neurons were identified, and AT8 fluorescence intensity was quantified on a per-neuron basis. AT8 signal was normalized to neuronal area and expressed as mean AT8 intensity per MAP2^+^ neuron. Data were normalized to control conditions and are reported as mean ± SEM from three independent experiments.

## AUTHOR INFORMATION

### Corresponding Author

Moustafa T. Gabr - Department of Radiology, Molecular Imaging Innovations Institute (MI3), Weill Cornell Medicine, New York, NY 10065, United States;

## ASSOCIATED CONTENT

The following files are available free of charge. LC-MS data of the synthesized peptides (PDF).

### Notes

The authors declare no competing financial interest.

## Insert Table of Contents artwork here

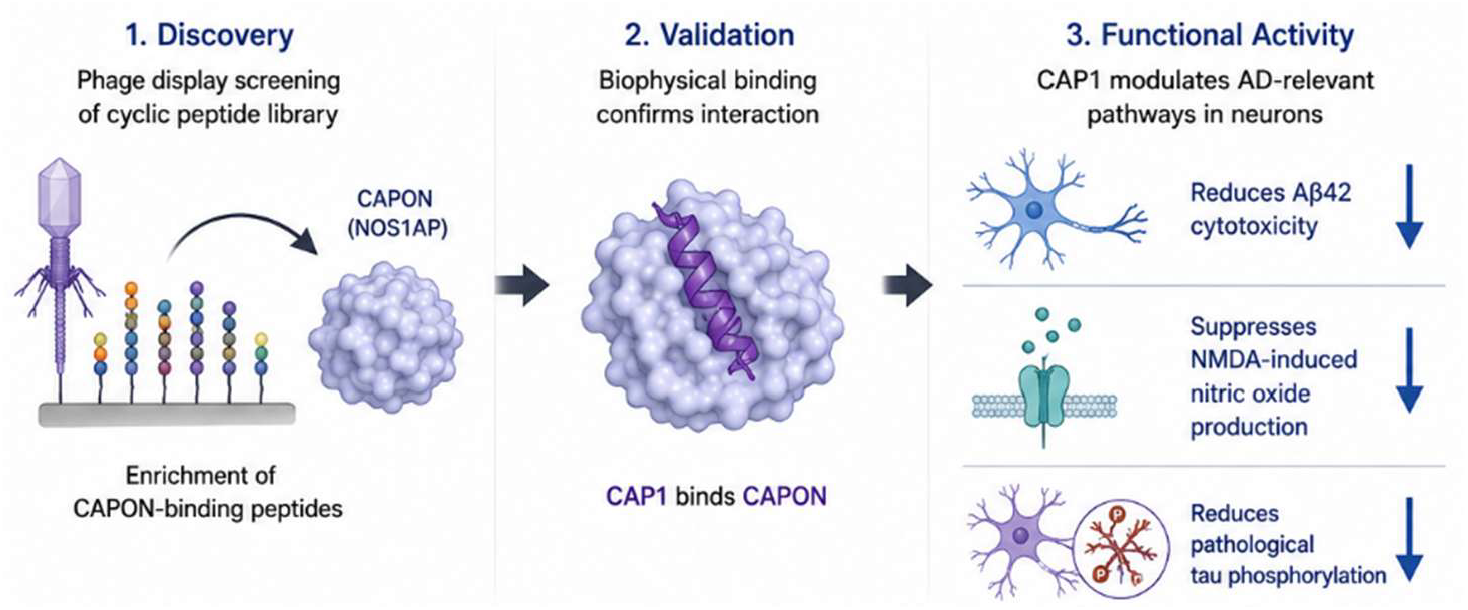

