## Supporting Information for "Cyclic Peptides Target CAPON and Modulate Cellular Responses under Alzheimer’s Disease-Relevant Stress"

*Electronic Supplementary Material for*

Ashraf Abdo,^1^ Shaoren Yuan,^1^ Katarzyna Kuncewicz,^1,2^ Jisong Mo,^3^ [Hongliang Duan](https://www.sciencedirect.com/author/57213838871/hongliang-duan),^3^ Moustafa T. Gabr^1*^

^1^Department of Radiology, Molecular Imaging Innovations Institute (MI3), Weill Cornell Medicine, New York, NY 10065, USA.

^2^Department of Biomedical Chemistry, Faculty of Chemistry, University of Gdansk, Poland

^3^ Faculty of Applied Sciences, Macao Polytechnic University, Macao 999078, China

Table of Contents

LC-MS spectra for CAP1	S2

LC-MS spectra for CAP2	S3

LC-MS spectra for CAP3	S4

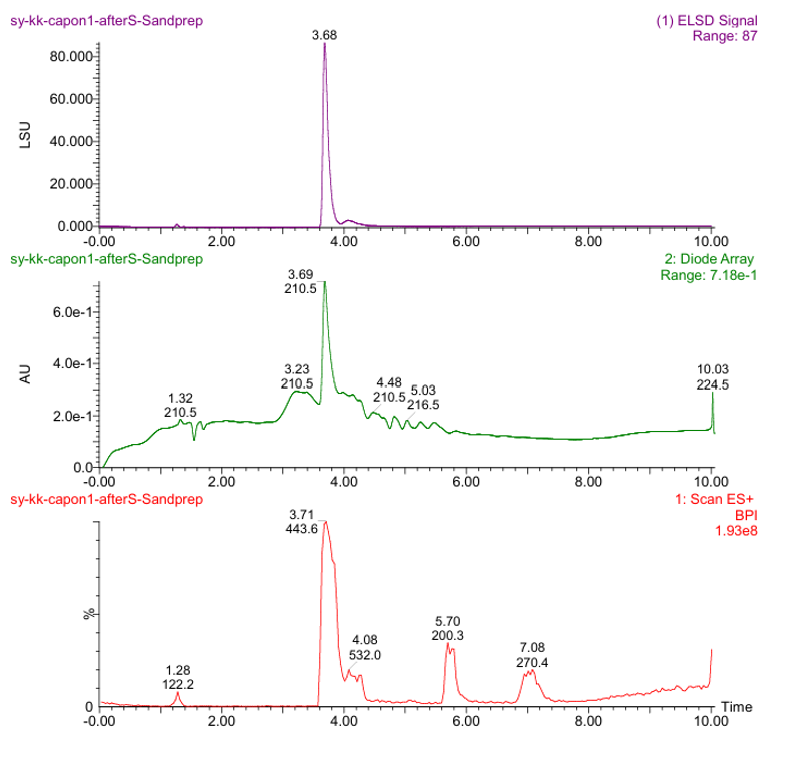

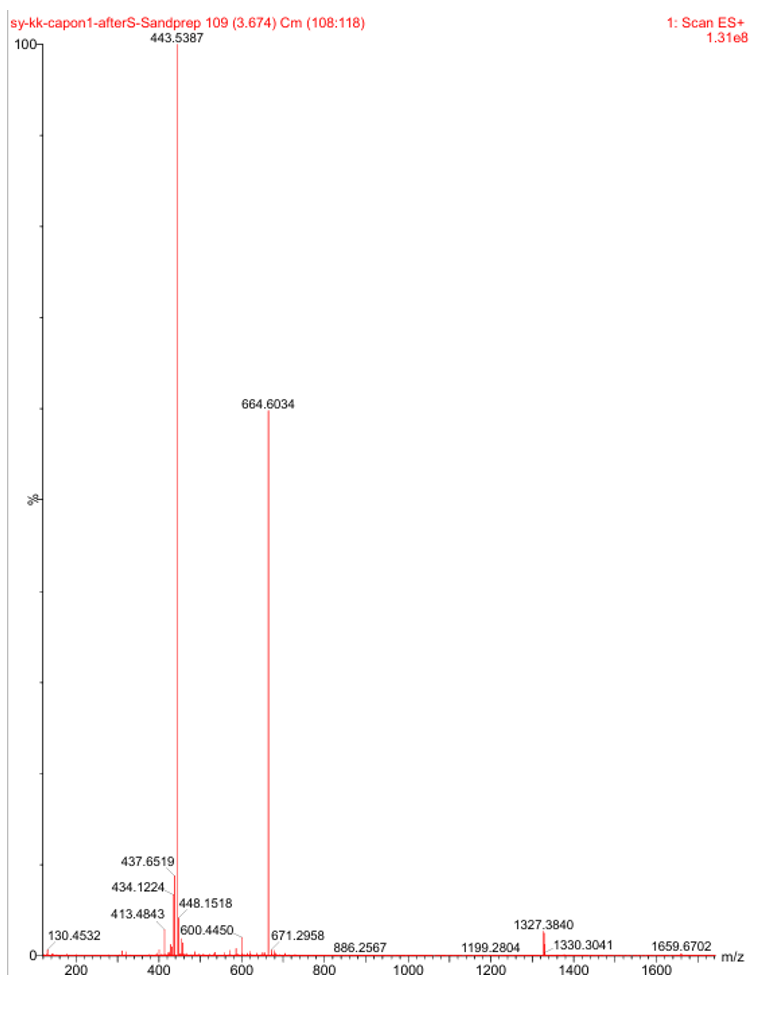

**Figure S1.** **LC-MS spectra for CAP1.**

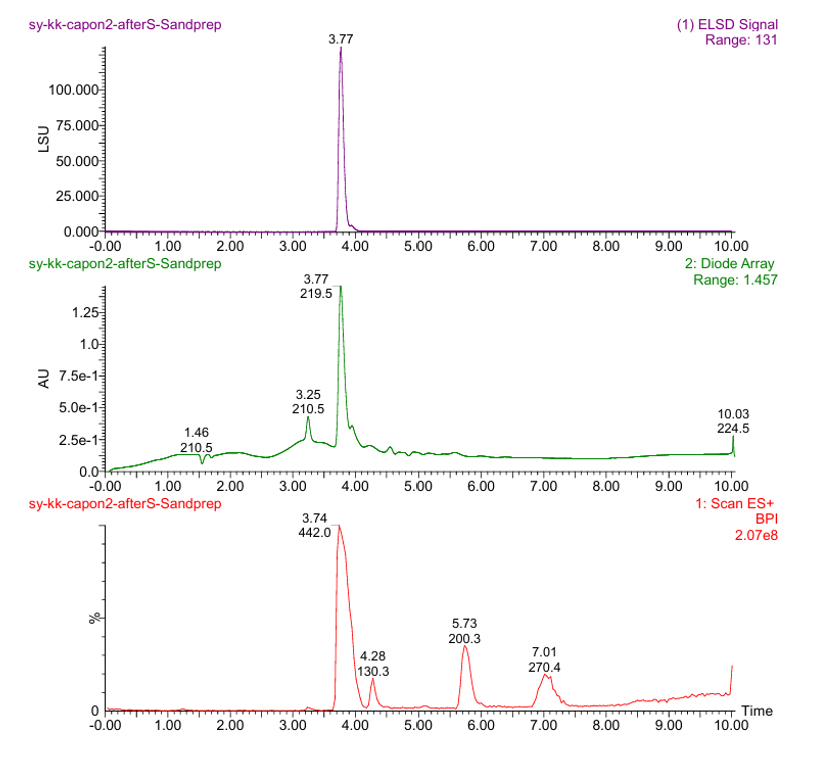

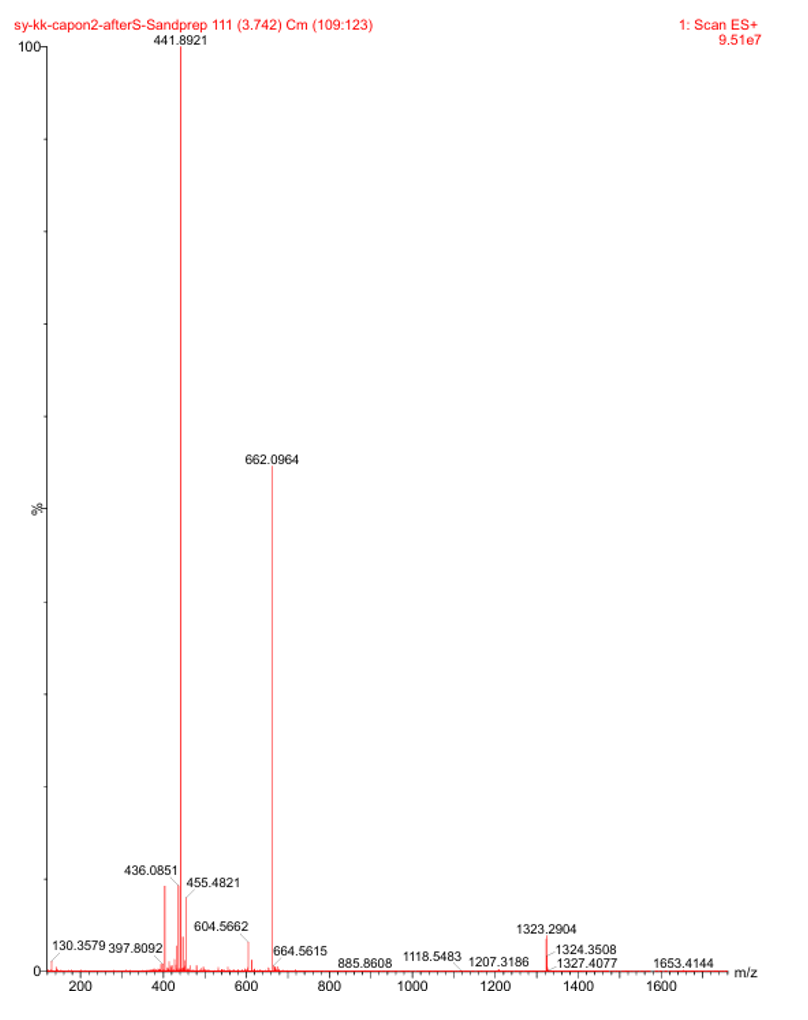

**Figure S2.** **LC-MS spectra for CAP2.**

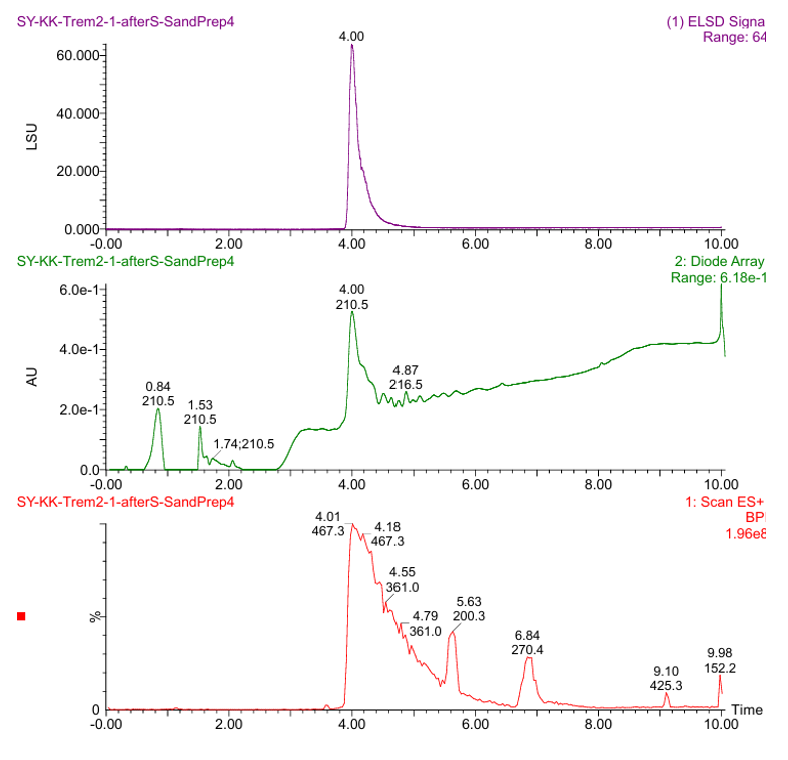

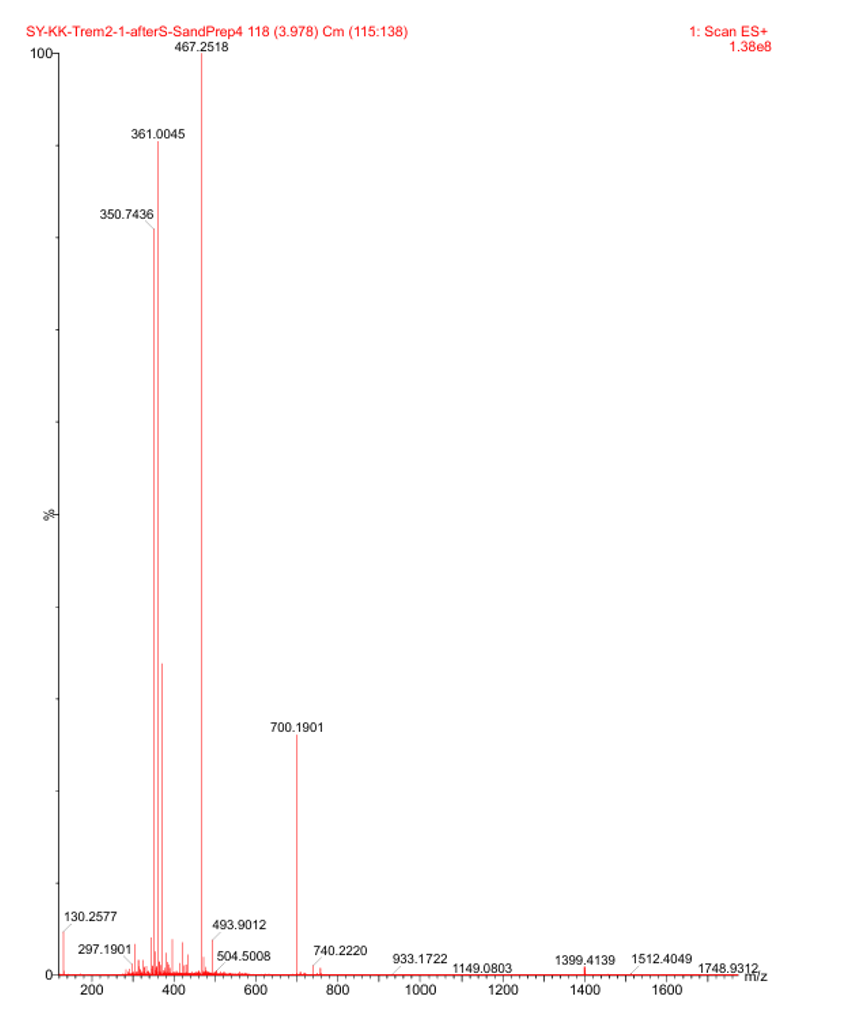

**Figure S3.** **LC-MS spectra for CAP3.**
